# Structure-Guided Design of C5aR1-Selective Peptide Agonists

**DOI:** 10.64898/2026.09.02.746116

**Authors:** Joshua Dent, Xiaosa Wu, Liam Rogl, Richard Clark

## Abstract

Selective peptide agonists for complement C5a receptor 1 (C5aR1) are valuable tools for dissecting receptor-specific inflammatory signalling, but their optimisation is complicated by overlap with closely related anaphylatoxin receptors and by pathway-dependent pharmacology. Here, we applied a structure-guided computational workflow to prioritise mutations within two C5a-derived peptide agonist scaffolds. FoldX-guided modelling identified position 5 as a candidate optimisation site, with hydrophobic substitutions predicted to improve C5aR1 engagement without corresponding gains at C3aR. Predicted BM1 and BM221 analogues were synthesised by solid-phase peptide synthesis and evaluated across C3aR, C5aR1 and C5aR2 using ERK1/2 phosphorylation and β-arrestin recruitment assays. Position-5 substitutions enhanced C5aR1 functional preference in ERK assays, although additional replacement of Leu^6^ with Ala reduced target potency and revealed pathway-dependent receptor discrimination. BM1 P^5^M provided the clearest overall improvement across ERK and β-arrestin readouts. In the BM221 series, A^5^Nle improved C5aR1 preference over C3aR, whereas A^5^Nle Abu^6^Ala produced the most favourable serum stability profile. These findings support position 5 as a transferable optimisation site and demonstrate that C5aR1 potency and receptor selectivity must be balanced during next-generation agonist design.

**Table of Contents Graphic:** 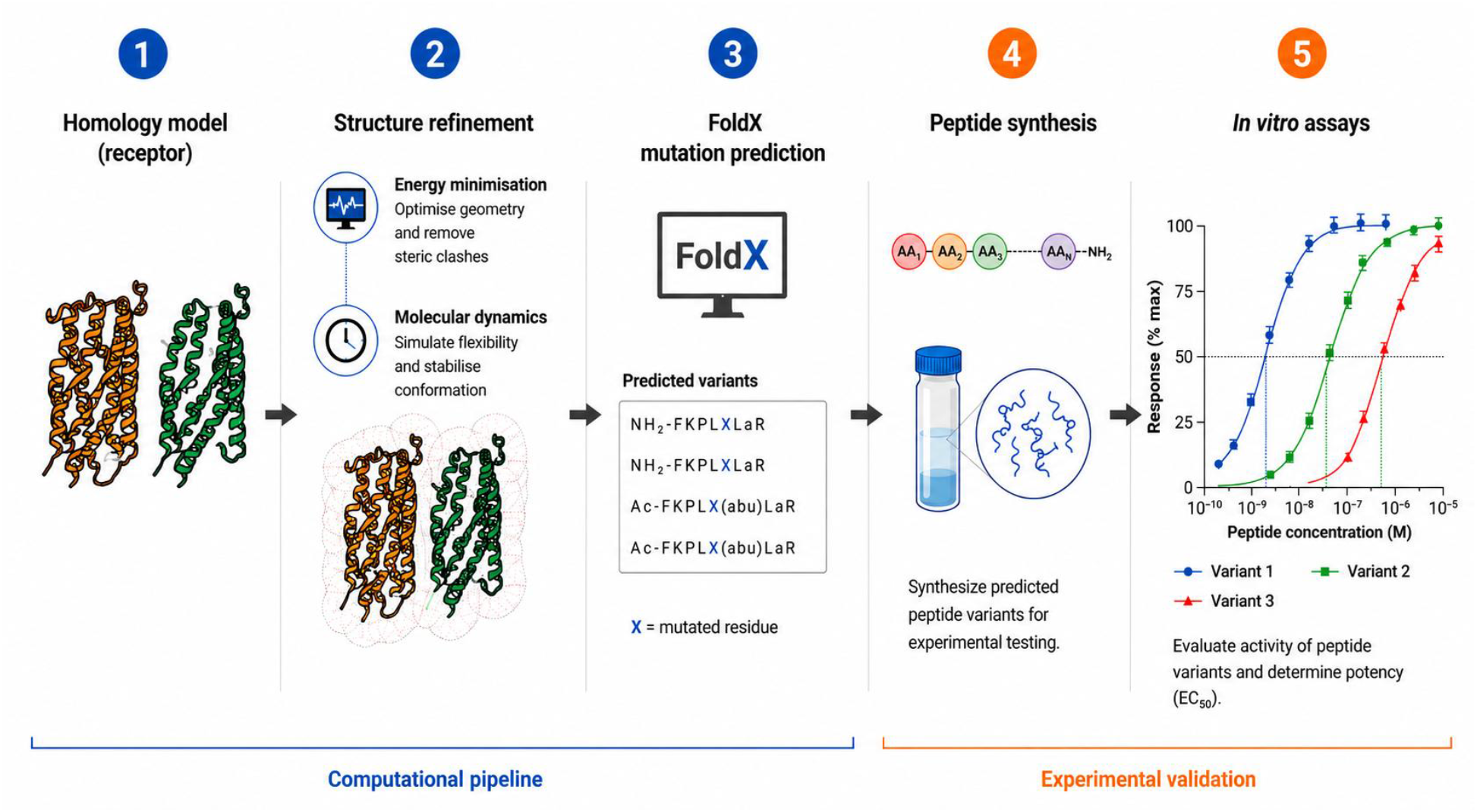

## Introduction

G protein-coupled receptors (GPCRs) regulate diverse physiological processes, including neurotransmission, metabolism and immune responses. Ligand binding stabilises receptor conformations that promote coupling to heterotrimeric G proteins and downstream effector pathways, while receptor phosphorylation by GPCR kinases supports β-arrestin recruitment, receptor desensitisation and internalisation^1-3^. β-arrestins can also act as signalling scaffolds, enabling G protein-independent activation of pathways such as ERK1/2 and JNK3^4, 5^. Because individual ligands can stabilise distinct active receptor conformations, GPCR signalling is often pathway-selective rather than uniform across all outputs^6^. This ligand-directed signalling complexity is particularly relevant for peptide–GPCR systems, where small sequence changes can alter potency, efficacy, receptor subtype selectivity and signalling bias^7, 8^.

C5aR1 is a prototypical immune GPCR activated by the complement anaphylatoxin C5a. C5aR1 couples predominantly to Gi/Go proteins and recruits β-arrestins, thereby supporting both second messenger signalling and receptor trafficking^9-11^. Receptor activation drives inflammatory processes including chemotaxis, degranulation, intracellular Ca^2+^ mobilisation and superoxide production, consistent with its central role in neutrophil and macrophage recruitment during acute inflammation^12-16^. Dysregulated C5a–C5aR1 signalling has also been implicated across chronic inflammatory and neurodegenerative disease contexts, including inflammatory arthritis, inflammatory bowel disease and Alzheimer’s disease^17-22^. Although C5aR1 inhibition has been the dominant therapeutic strategy, selective agonists provide complementary tools for dissecting receptor activation mechanisms, pathway bias and cell-type-specific complement signalling. Prior C5aR1-selective agonists have already demonstrated that receptor-selective activation can produce distinct signalling profiles and measurable in vivo activity, supporting their use as pharmacological probes^7, 23, 24^. Accordingly, C5aR1-directed ligands must be optimised not only for potency, but also for receptor selectivity and pathway engagement.

C5a recognition by C5aR1 is commonly described as a multi-site interaction. The receptor N-terminus contributes to high-affinity ligand engagement, whereas the C-terminal region of C5a interacts with the transmembrane core to promote receptor activation. A further interaction between extracellular loop 2 and a hydrophobic region of C5a helps stabilise the receptor–ligand complex^25-27^. These features have motivated the development of short C5a-derived peptide agonists that retain the minimal activation motif of the endogenous ligand^28-30^. However, C5a-derived peptides do not necessarily reproduce the full pharmacological profile of C5a, with some retaining robust proximal or ERK-linked signalling while showing reduced β-arrestin recruitment or receptor internalisation^11, 24^. Thus, optimisation of C5aR1 peptide agonists requires simultaneous consideration of receptor subtype selectivity and pathway engagement.

C5a also binds with similar potency to a second receptor, C5a receptor 2 (C5aR2), which is co-expressed with C5aR1 across a wide range of both myeloid and non-myeloid cell types^31-34^. C5aR2 has not been shown to signal through G proteins, a finding initially attributed to divergence in the DRY and NPxxY motifs, as well as differences in intracellular loop architecture^35-37^. As C5a is a cognate ligand for both C5aR1 and C5aR2, C5a-derived peptide agonists based on the minimal C-terminal motif may also retain activity at both receptor subtypes. Thus, C5aR2 activity among shortened C5a mimetics is not unexpected, but reflects the native dual-receptor pharmacology of C5a. By contrast, C3aR activity suggests that isolating the C5a C-terminal pharmacophore from its native scaffold can reduce receptor discrimination across the broader anaphylatoxin receptor family. This issue of receptor promiscuity is evident among several C5a-derived ligands such as EP54, EP67 and C5apep^11, 38^. These examples demonstrate that apparent C5aR1 potency must be interpreted alongside counter-screening at related anaphylatoxin receptors.

The BM-series of peptides provides an experimentally validated starting point for C5aR1 agonist optimisation. Within this series, BM1 serves as a useful parent scaffold because it retains robust C5aR1 activity while exhibiting limited receptor discrimination, providing a baseline for assessing improvements in receptor selectivity. Previous work further identified BM213 as a C5aR1 agonist biased against β-arrestin recruitment and BM221 as a C5aR1-selective agonist without detectable signalling bias under the conditions tested^7^. Together, these peptides provide complementary pharmacological benchmarks: BM1 enables assessment of whether computationally prioritised mutations can improve selectivity from a less selective starting point, whereas BM221 allows the same mutations to be tested within an already C5aR1-selective scaffold. Their synthetic accessibility and prior characterisation make them well suited for testing how targeted sequence changes affect receptor preference, pathway engagement and peptide developability.

Peptide ligand optimisation has traditionally relied on empirical screening and iterative structure– activity relationship analysis^39^. Although these approaches have been productive, they can become inefficient when receptor selectivity and pathway-dependent pharmacology cannot be predicted from sequence information alone. Recent advances in GPCR structural biology, particularly the determination of active-state cryo-EM receptor–transducer complexes, have enabled more rational structure-based design of peptide ligands. Static structures nevertheless provide only limited snapshots of receptor conformational landscapes^40-42^. Molecular dynamics simulations and empirical energy-based tools such as FoldX can complement structural models by estimating the effects of peptide substitutions on receptor–ligand interactions and by prioritising experimentally testable analogues^43-45^.

A precedent for this approach was provided by our previous computational design of α-conotoxins with improved selectivity among nicotinic acetylcholine receptor subtypes^46^. In that work, experimentally informed peptide–receptor complex models were refined and subjected to FoldX-based mutational energy calculations, allowing a focused set of substitutions to be prioritised for synthesis and functional evaluation. This workflow provided a practical means of directing experimental SAR toward high-value variants without requiring extensive empirical peptide libraries. The availability of published C5aR1–BM213 and C3aR–EP54 structures provided an opportunity to adapt this model-building, mutational scanning and experimental validation strategy to the BM peptide series.

Here, C5aR1–BM peptide and C3aR–BM peptide models were generated from the available experimental structures and evaluated against existing activity data before FoldX-based mutational calculations were used to prioritise substitutions at selected peptide positions. BM1 and BM221 were used as complementary peptide scaffolds to test the transferability and sequence dependence of the computationally prioritised substitutions. Selected analogues were synthesised by solid-phase peptide synthesis, purified by HPLC and evaluated at C3aR, C5aR1 and C5aR2 using ERK1/2 phosphorylation and β-arrestin recruitment assays. Selected BM221 analogues were additionally assessed for stability in human serum. Collectively, this study examined whether structure-guided computational prioritisation could identify C5aR1 agonists that retain robust target activity while improving functional receptor discrimination and, where applicable, peptide stability.

## Results

### Computational prioritisation identified position 5 as a candidate C5aR1 optimisation site

Changes in predicted binding free energy caused by amino acid substitutions in the BM1 scaffold were evaluated using homology modelling, molecular dynamics refinement and FoldX calculations. The resulting ΔΔG profiles identified position 5 as a candidate site for improving C5aR1 engagement. Replacement of Pro^5^ with hydrophobic residues was predicted to be favourable in the C5aR1 model, with P^5^M and P^5^L producing ΔΔG values of −1.42 and −1.19 kcal/mol, respectively. In contrast, equivalent substitutions were not predicted to produce comparable gains in the C3aR model. These predictions suggested that linear hydrophobic side chains at position 5 may improve C5aR1 interactions while avoiding parallel increases in C3aR engagement.

On this basis, Pro^5^-to-Met and Pro^5^-to-Leu substitutions were selected in the BM1 scaffold. In addition to these computationally prioritised analogues, an L^6^A substitution was introduced into Ac-BM1 P5L based on the structure–activity relationship previously reported for BM213. BM213 contains Ala at position 6 and retains C5aR1-mediated calcium mobilisation and ERK1/2 phosphorylation while showing no detectable β-arrestin recruitment.^7^ This distinctive signalling profile suggested that reduced side-chain bulk at position 6 may contribute to pathway-dependent C5aR1 activation. Accordingly, Ac-BM1 P^5^L L^6^A was designed to determine whether this previously identified position-6 feature was transferable to the P5L scaffold. Corresponding position-5 modifications were also introduced into BM221 to test whether the predicted optimisation site was transferable across related C5aR1 agonist scaffolds. The BM221 series included A^5^M, A^5^Nle and A^5^Nle Abu^6^Ala analogues, enabling assessment of hydrophobic side-chain length and the contribution of the adjacent Abu6 position to functional activity and serum stability

**Figure 2:**
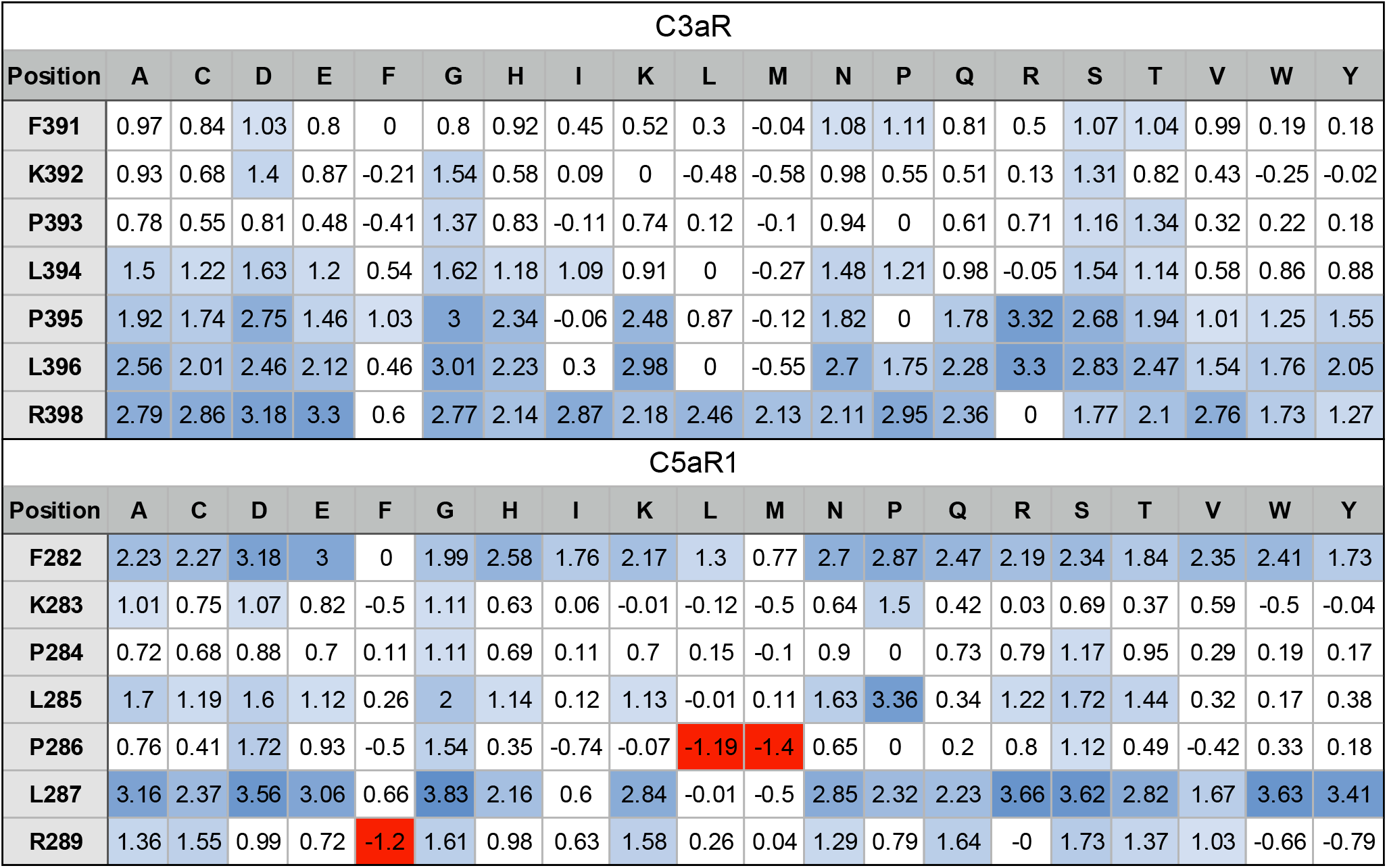
Changes in Binding Free Energies of BM1 analogues to C3aR and C5aR1. Predicted changes to free binding energies of BM1 mutants binding to C3aR (**Top**) and C5aR1 (**Bottom**). Δ ΔG values < -1 kcal/mol indicate an increase in binding affinity, whilst values of -1 < Δ ΔG < 1 kcal/mol indicate no change in binding affinity. Finally Δ ΔG > 1 kcal/mol indicates a reduced affinity for the receptor. BM1 analogues with leucine or methionine substitutions at position 5 were predicted to have increased affinity (shown in red) for C5aR1 with no corresponding increase at C3aR.

### Position-5 substitutions improved C5aR1 functional preference in the BM1 scaffold

**Table 1.** Pharmacological activity of BM1 position-5 analogues.

| Peptide | Sequence | pERK 1/2 pEC<sub>50</sub> ± SE<sup>a</sup> |  | β-arrestin pEC<sub>50</sub> ± SE<sup>b</sup> |  |
| --- | --- | --- | --- | --- | --- |
|  |  | C5aR1 | C3aR | C5aR1 | C5aR2 |
| BM1 | NH2-FKPLPLaR | 7.82 (0.265) | 6.33 (0.152) | 5.88 (0.109) | 4.28 (0.082) |
| BM213 | Ac-FKPLAAaR | 7.23 (0.115) | 4.26 (0.096) | ND | ND |
| BM221 | Ac-FKPLA(Abu)aR | 7.48 (0.218) | 4.77 (0.329) | 5.49 (0.068) | 5.15 (0.174) |
| P<sup>5</sup>M | NH2-FKPLMLaR | 9.01 (0.479) | 5.54 (0.351) | 6.98 (0.128) | 4.23 (0.067) |
| P<sup>5</sup>L | NH2-FKPLLLaR | 8.96 (0.349) | 4.94 (0.199) | 5.97 (0.154) | 4.09 (0.067) |
| Ac-P<sup>5</sup>L | Ac-FKPLLLaR | 8.69 (0.227) | 6.44 (0.134) | 5.49 (0.051) | 5.03 (0.362) |
| Ac-P<sup>5</sup>L L<sup>6</sup>A | Ac-FKPLLAaR | 7.03 (0.125) | 4.29 (0.245) | 5.09 (0.066) | 5.21 (0.218) |

<sup>a</sup>pERK 1/2 signalling was measured in CHO cells stably expressing C5aR1 or C3aR. <sup>b</sup>β-Arrestin recruitment was measured using BRET-based assays in transiently transfected HEK293 cells. N = 3–6 unless otherwise indicated. ND = not determined. Ac- indicates N-terminal acetylation.

The BM1 position-5 analogues were first assessed for receptor activity across C3aR and C5aR1 in ERK1/2 phosphorylation assays. Consistent with the computational design objective, BM1 P^5^L and BM1 P^5^M showed reduced C3aR potency relative to BM1, while each analogue retained robust C5aR1-mediated ERK activation. In the C5aR1 pERK assay, both BM1 P^5^M and BM1 P^5^L were left-shifted relative to BM1, indicating increased potency at the target receptor. Ac-BM1 P^5^L also retained potent C5aR1 activity, although N-terminal acetylation altered its off-target receptor profile relative to the non-acetylated P5L analogue. Replacement of the adjacent Leu^6^ residue with Ala in Ac-BM1 P5L L6A reduced C5aR1 potency relative to Ac-BM1 P5L but produced a larger reduction in C3aR potency, thereby maintaining a preference for C5aR1 in the ERK readout.

β-Arrestin recruitment revealed more pathway-dependent effects of the BM1 modifications. BM1 P^5^M produced the clearest gain in C5aR1 β-arrestin potency relative to BM1, whereas BM1 P^5^L and Ac-BM1 P^5^L remained closer to the parent scaffold. At C5aR2, BM1, BM1 P^5^M and BM1 P^5^L remained largely inactive until high concentrations, whereas Ac-BM1 P^5^L showed greater C5aR2 engagement. Ac-BM1 P^5^L L^6^A displayed lower C5aR1 β-arrestin potency than Ac-BM1 P^5^L and retained measurable C5aR2 activity, resulting in little apparent discrimination between C5aR1 and C5aR2. Selectivity analysis using ΔpEC50 values further supported the conclusion that position-5 modification improved C5aR1 preference over C3aR in the ERK readout, with BM1 P^5^M providing the most balanced improvement across ERK and β-arrestin assays.

**Figure 3.**
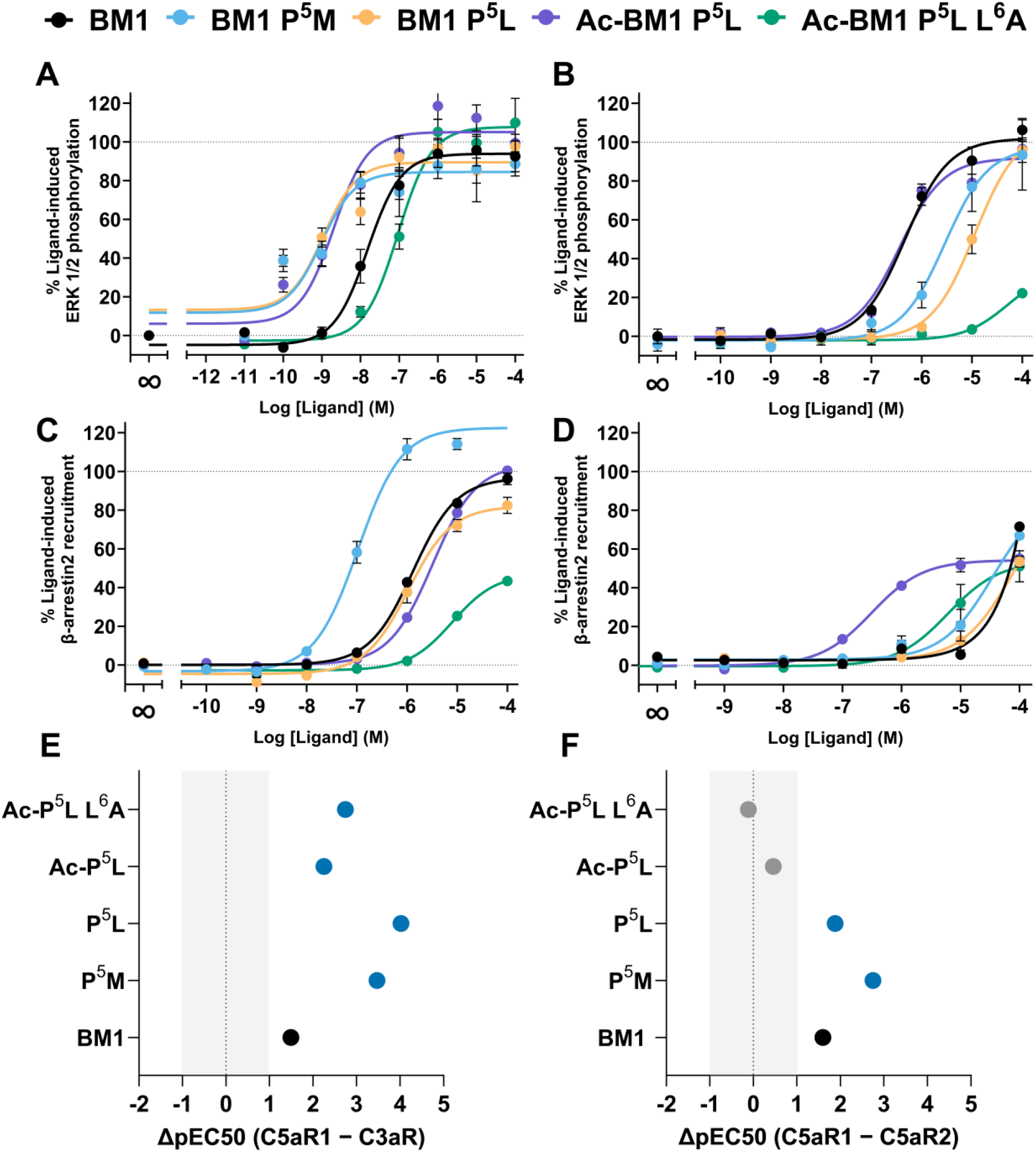
Pharmacological profiling of BM1 analogues at C5aR1, C3aR and C5aR2. (**A, B**) Concentration–response curves for ligand-induced ERK1/2 phosphorylation in CHO cells stably expressing C5aR1 (A) or C3aR (B), measured using the AlphaLISA SureFire Ultra p-ERK1/2 assay. Responses were normalised to vehicle-treated wells (0%) and the corresponding endogenous agonist control (100%; 100 nM C5a for C5aR1 or 100 nM C3a for C3aR). (**C, D**) Concentration–response curves for ligand-induced β-arrestin 2 recruitment in HEK293 cells transiently expressing C5aR1-Rluc8/β-arrestin 2-Venus (C) or C5aR2-Venus/β-arrestin 2-Rluc8 (D). Responses were normalised to vehicle-treated wells (0%) and 100 nM C5a-treated wells (100%). (**E, F**) Receptor preference was summarised from potency differences between C5aR1 and competing receptors for ERK1/2 phosphorylation (E; C3aR versus C5aR1) or β-arrestin 2 recruitment (F; C5aR2 versus C5aR1). Positive values indicate greater apparent potency at C5aR1 relative to the competing receptor. The shaded region denotes values within ±1 log unit, corresponding to less than 10-fold preference. Blue points indicate >10-fold preference for C5aR1, grey points indicate little or no receptor preference. Data are presented as mean ± SEM from N = 3–4 independent experiments for panels A–C and N = 3 independent experiments for panel D, with each concentration tested in technical triplicate. Panels E and F show mean ΔpEC50 values from three independent experiments.

### Position-5 modification was transferable to the BM221 scaffold but did not improve C5aR1:C5aR2 discrimination

To determine whether the position-5 optimisation strategy was scaffold-dependent, analogous substitutions were introduced into the BM221 scaffold. In C3aR pERK assays, BM221 and the position-5 analogues showed limited activation overall, with responses generally emerging only at higher concentrations. In contrast, all BM221-derived peptides acted as full agonists in the C5aR1 pERK assay relative to the assay control, indicating that position-5 modification preserved robust C5aR1 ERK signalling. BM221 A^5^M and BM221 A^5^Nle showed increased C5aR1 potency relative to BM221, consistent with the predicted benefit of hydrophobic substitution at this position.

In C5aR1 β-arrestin2 recruitment assays, BM221 A5M and BM221 A5Nle showed increased apparent potency relative to BM221, whereas BM221 A5Nle Abu6Ala showed reduced apparent potency. All three modified analogues retained measurable C5aR2 activity. BM221 A5M and BM221 A5Nle Abu6Ala each showed modest apparent C5aR1 preference, with ΔpEC50 values of 0.66, while BM221 A5Nle showed essentially no receptor preference. Thus, none of the BM221 analogues achieved greater than 10-fold functional discrimination between C5aR1 and C5aR2.

**Figure 4.**
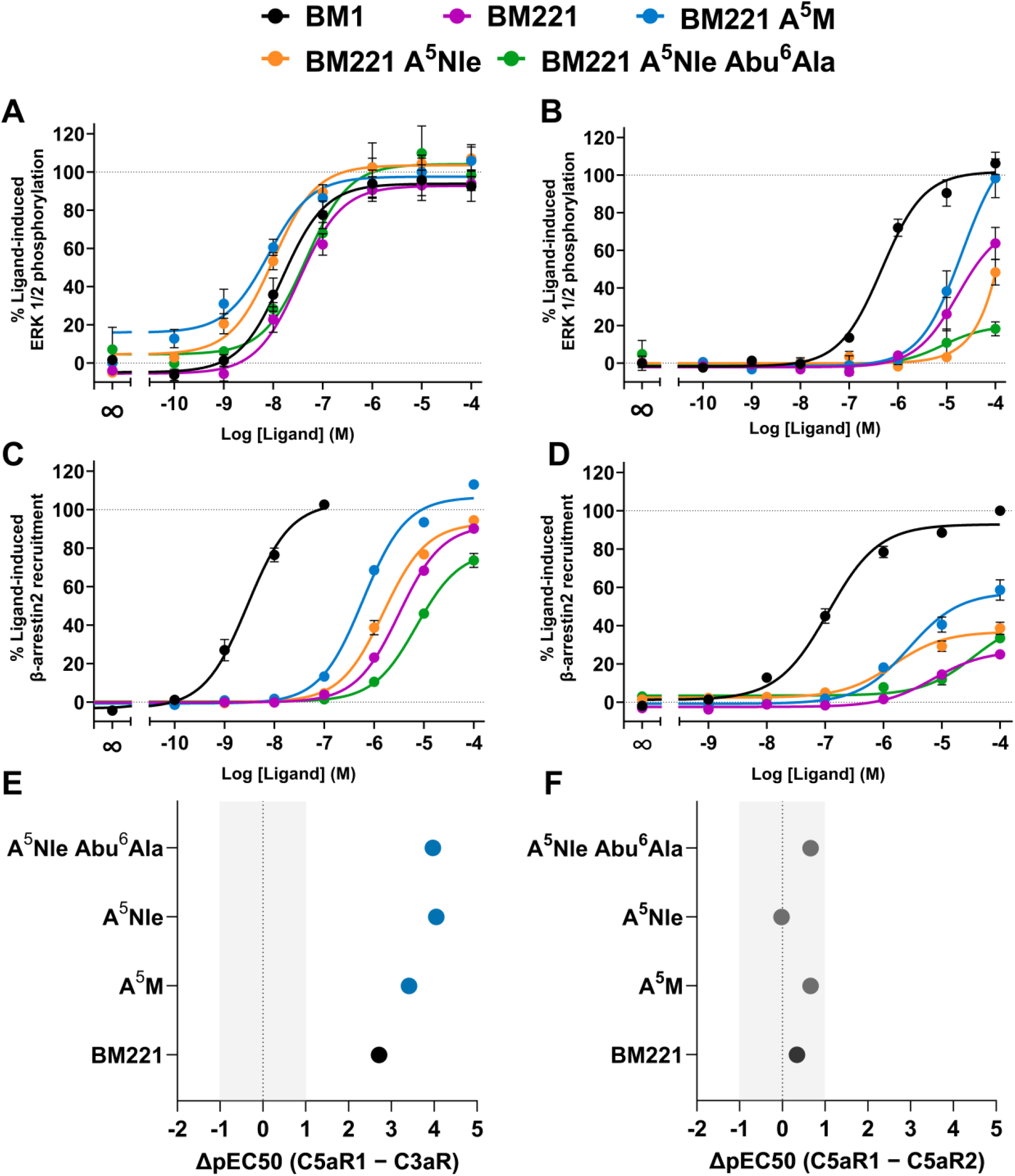
Pharmacological profiling of BM221 analogues at C5aR1, C3aR and C5aR2. (**A, B**) Concentration–response curves for ligand-induced ERK1/2 phosphorylation in CHO cells stably expressing C5aR1 (A) or C3aR (B), measured using the AlphaLISA SureFire Ultra p-ERK1/2 assay. Responses were normalised to vehicle-treated wells (0%) and the corresponding endogenous agonist control (100%; 100 nM C5a for C5aR1 or 100 nM C3a for C3aR). (**C, D**) Concentration–response curves for ligand-induced β-arrestin 2 recruitment in HEK293 cells transiently expressing C5aR1-Rluc8/β-arrestin 2-Venus (C) or C5aR2-Venus/β-arrestin 2-Rluc8 (D). Responses were normalised to vehicle-treated wells (0%) and 100 nM C5a-treated wells (100%). (**E, F**) Receptor preference was summarised from potency differences between C5aR1 and comparator receptors for ERK1/2 phosphorylation (E; C3aR versus C5aR1) or β-arrestin 2 recruitment (F; C5aR2 versus C5aR1). Positive values indicate greater apparent potency at C5aR1 relative to the comparator receptor. The shaded region denotes values within ±1 log unit, corresponding to less than 10-fold preference. Blue points indicate >10-fold preference for C5aR1, grey points indicate little or no receptor preference, and orange points, where present, indicate >10-fold preference for the comparator receptor. Data are presented as mean ± SEM from N = 3–4 independent experiments for panels A–C and N = 3 independent experiments for panel D, with each concentration tested in technical triplicate. Panels E and F show mean ΔpEC50 values from three independent experiments.

### BM221 A5Nle Abu6Ala showed the greatest serum stability among the tested analogues

**Table 2.**
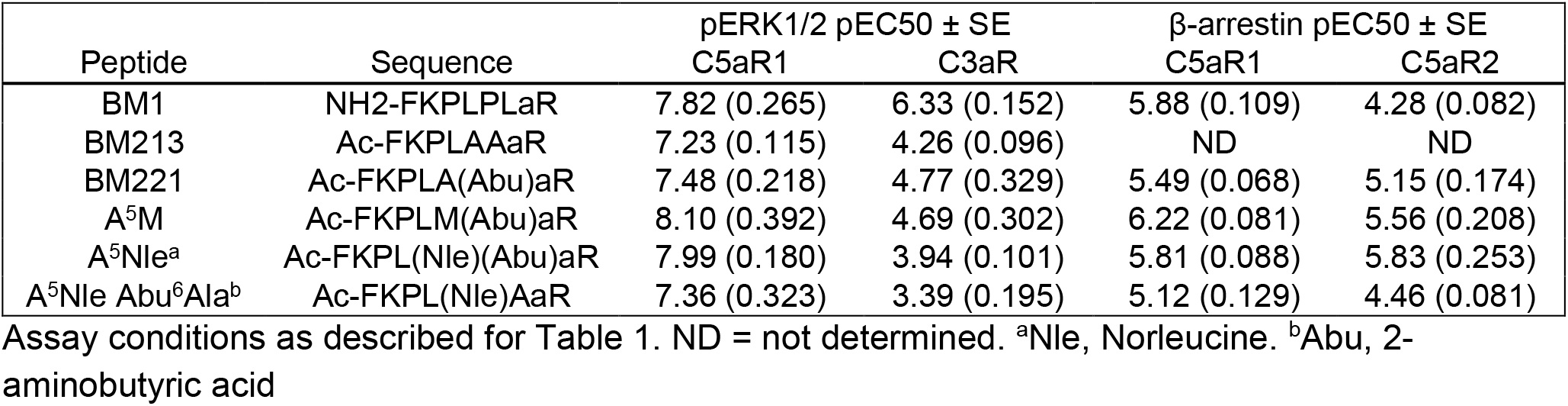
Pharmacological activity of BM221 position-5 analogues.

| Peptide | Sequence | pERK1/2 pEC<sub>50</sub> ± SE | | β-arrestin pEC<sub>50</sub> ± SE | |
| --- | --- | --- | --- | --- | --- |
|  |  | C5aR1 | C3aR | C5aR1 | C5aR2 |
| BM1 | NH<sub>2</sub>-FKPLPLaR | 7.82 (0.265) | 6.33 (0.152) | 5.88 (0.109) | 4.28 (0.082) |
| BM213 | Ac-FKPLAAaR | 7.23 (0.115) | 4.26 (0.096) | ND | ND |
| BM221 | Ac-FKPLA(Abu)aR | 7.48 (0.218) | 4.77 (0.329) | 5.49 (0.068) | 5.15 (0.174) |
| A<sup>5</sup>M | Ac-FKPLM(Abu)aR | 8.10 (0.392) | 4.69 (0.302) | 6.22 (0.081) | 5.56 (0.208) |
| A<sup>5</sup>Nle<sup>a</sup> | Ac-FKPL(Nle)(Abu)aR | 7.99 (0.180) | 3.94 (0.101) | 5.81 (0.088) | 5.83 (0.253) |
| A<sup>5</sup>Nle Abu<sup>6</sup>Ala<sup>b</sup> | Ac-FKPL(Nle)AaR | 7.36 (0.323) | 3.39 (0.195) | 5.12 (0.129) | 4.46 (0.081) |

Assay conditions as described for Table 1. ND = not determined. <sup>a</sup>Nle, Norleucine. <sup>b</sup>Abu, 2-aminobutyric acid

Because receptor selectivity alone does not determine peptide utility, selected BM221 analogues were evaluated for stability in 100% human serum using RP-HPLC. Intact peptide was quantified by integrated peak area and normalised to the corresponding t = 0 time point. BM221 A^5^Nle was less stable than BM221, with a fitted half-life of 3.1 h compared with 4.3 h for BM221. In contrast, BM221 A^5^Nle Abu^6^Ala displayed the greatest serum stability, with a fitted half-life of 8.1 h. This represented an approximately two-fold increase relative to BM221 and an approximately three-fold increase relative to BM221 A^5^Nle. Comparison of the serum degradation curves by nonlinear regression using an extra sum-of-squares F test indicated that the peptides exhibited significantly different degradation profiles (P < 0.0001), with BM221 A^5^Nle Abu^6^Ala showing the slowest rate of degradation.

These data show that hydrophobic substitution at position 5 does not uniformly improve serum stability. Instead, the increased stability of BM221 A^5^Nle Abu^6^Ala suggests that developability gains depend on the combined sequence context rather than on position-5 hydrophobicity alone. Thus, BM221 A5Nle Abu6Ala showed the greatest serum stability among the tested peptides despite its reduced C5aR1 β-arrestin2 potency and less than 10-fold functional preference for C5aR1 over C5aR2.

**Figure 5.**
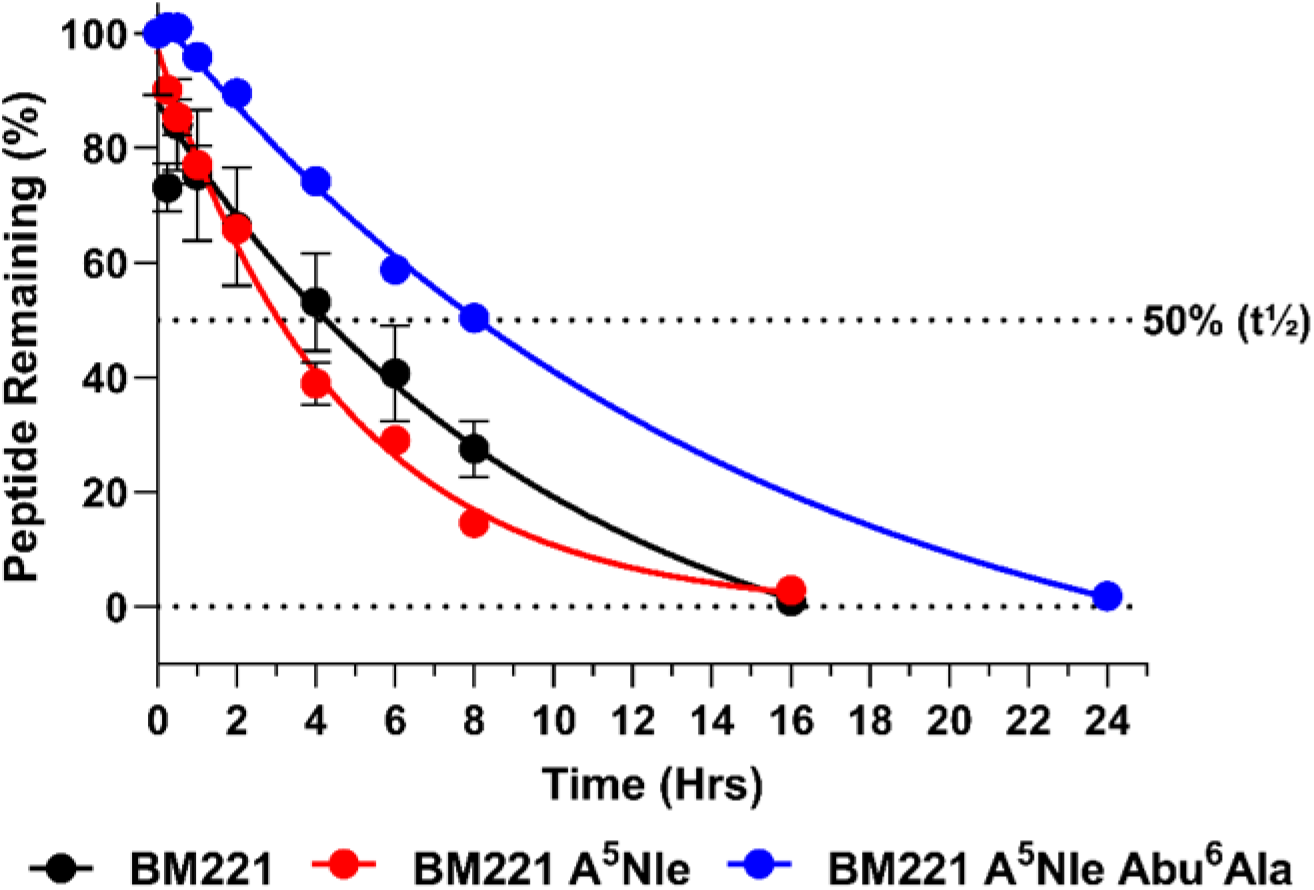
Serum stability of BM221 and optimised analogues in human serum. Peptides were incubated in 100% human serum at 37 °C, and the percentage of intact peptide remaining was quantified by RP-HPLC relative to the corresponding t = 0 sample. Data are presented as mean ± SEM from three independent experiments. Sequence optimisation differentially affected serum stability, with half-lives of 4.3 h (BM221), 3.1 h (BM221 A^5^Nle), and 8.1 h (BM221 A^5^Nle Abu^6^Ala). The dashed line indicates 50% peptide remaining.

## Discussion and Conclusions

This study demonstrates that structure-guided computational design can complement the empirical SAR approaches used to develop short C5aR1 agonists^46^. Computational prioritisation identified position 5 as a candidate optimisation site across the BM1 and BM221 scaffolds, and experimental evaluation confirmed that hydrophobic substitution at this position could alter C5aR1 potency and functional discrimination from related anaphylatoxin receptors. However, the effects depended on scaffold and adjacent sequence context, and improvements in receptor preference, pathway engagement and serum stability did not consistently coincide. These findings establish position 5 as a useful design site while reinforcing the need for experimental counter-screening across receptors and signalling pathways.

The BM1 series provided the clearest experimental support for the computational prioritisation of position 5. Position-5 substitution reduced functional activity at C3aR while preserving or increasing activity at C5aR1, consistent with a selectivity-by-disruption model in which the substitutions are less compatible with productive C3aR engagement or activation while remaining tolerated within C5aR1. Ac-BM1 P^5^M showed the most balanced profile, improving C5aR1 ERK potency and β-arrestin recruitment without the detectable C5aR2 β-arrestin recruitment observed for Ac-BM1 P^5^L. Position-5 modification therefore altered both receptor preference and pathway output rather than acting as a non-specific potency enhancer. Introduction of L^6^A into Ac-BM1 P^5^L reduced C5aR1 β-arrestin potency but also substantially reduced ERK potency and did not eliminate C5aR2 recruitment. The inability of this substitution to reproduce the pharmacological profile of BM213 indicates that Ala^6^ does not independently determine pathway engagement, but acts in conjunction with the surrounding sequence, including the adjacent position 5 residue^7^.

The BM221 series tested whether this optimisation site could be transferred to a distinct C5aR1-selective scaffold^7^. BM221 A^5^M and BM221 A^5^Nle increased C5aR1 potency in ERK and β-arrestin assays, supporting position 5 as a generalisable SAR determinant. However, these analogues also retained measurable C5aR2 β-arrestin recruitment, highlighting an important constraint for next-generation C5aR1 agonist design. C5aR1 and C5aR2 share ligand-recognition features but differ in signalling behaviour. C5aR2 has not been shown to engage conventional G protein signalling but can recruit β-arrestins and modulate complement receptor responses^36, 47^. Consequently, β-arrestin recruitment provides a particularly important readout when counter-screening C5a-derived agonists at C5aR2. The retention of C5aR2 activity across the modified BM221 analogues indicates that changes in C5aR1 β-arrestin2 potency do not inherently confer functional discrimination against C5aR2. Counter-screening at C5aR2 should therefore be incorporated early in future design cycles, particularly when β-arrestin recruitment is used as a key optimisation endpoint.

Available C5aR1 structural data provide a plausible rationale for the observed effects of the position 5 substitutions. Cryo-EM structures show that the C-terminus of C5a, as well as peptide ligands derived from this region, adopt a hook-shaped conformation in the orthosteric binding site of C5aR1^24^. Hydrophobic residues in C5a-derived peptide agonists occupy a hydrophobic region of the orthosteric pocket associated with TM2 and TM3, and improved occupancy of this region may stabilise receptor conformations linked to activation^24^. In C5apep, the side chain of cyclohexylalanine occupies the hydrophobic face of the pocket, with earlier SAR studies of C5a-derived octapeptides similarly identifying that hydrophobic substitutions at positions 4-6 improved receptor affinity^24, 48^. This interaction was further dependent on side-chain geometry, with bulkier aromatic residues such as phenylalanine and tyrosine reducing binding in a manner consistent with steric restriction in the binding pocket^48^. The favourable activity of Met and Leu substitutions in BM1, and Met and Nle substitutions in BM221, is consistent with this structural framework. In each scaffold, replacing the parent position-5 residue with a larger aliphatic side chain may improve complementarity with the TM2/TM3-facing hydrophobic region, although the starting residues differ between BM1 and BM221.

The context-dependent effect of the L^6^A substitution can be interpreted in light of available C5aR1 structural and mutagenesis data. In the C5a-bound C5aR1 structure, ligand Leu^6^ extends into a hydrophobic cleft formed by TM2, ECL1 and TM3. By contrast, the shorter Ala^6^ side chain of BM213 projects toward the same region but makes fewer hydrophobic contacts with the receptor^24^. Receptor and ligand mutagenesis studies further implicated the interaction between ligand position 6 and the I^91^–W^102^–I^116^ region in β-arrestin recruitment^24^. Substitution of Ala^6^ in BM213 with Val, Leu, Ile, norvaline or Met increased β-arrestin recruitment relative to BM213, whereas Phe and homoproline did not produce the same effect^24^. This SAR suggests that productive engagement of the hydrophobic cleft requires sufficient side chain reach and complementarity but may be impaired by excessive bulk or unsuitable geometry. Accordingly, the reduced β-arrestin potency of Ac-BM1 P^5^L L^6^A relative to Ac-BM1 P5L is consistent with weaker engagement of this region by the smaller Ala6 side chain. However, the retention of measurable β-arrestin recruitment by Ac-BM1 P^5^L L^6^A, together with its broader divergence from the pharmacological profile of BM213, indicates that position 6 does not function as an independent determinant of pathway engagement^7^. Instead, its contribution appears to depend on the broader peptide sequence, including the identity of the adjacent position 5 residue. Because no structures of the analogues developed in the present study were determined in complex with C5aR1, this interpretation remains a structure-guided mechanistic hypothesis rather than direct evidence of their binding modes.

The serum stability data further emphasise that pharmacological and developability optimisation are related but separable objectives. BM221 A^5^Nle improved C5aR1 functional preference over C3aR in the ERK1/2 assay but showed reduced serum stability relative to BM221. By contrast, BM221 A^5^Nle Abu^6^Ala showed the greatest serum stability among the tested peptides despite its reduced C5aR1 β-arrestin2 potency and limited functional discrimination between C5aR1 and C5aR2. The increased half-life of BM221 A^5^Nle Abu^6^Ala suggests that local sequence context can reduce proteolytic susceptibility, potentially by altering backbone flexibility or disrupting preferred cleavage motifs^49^. However, serum stability should be considered an early developability measure rather than a direct proxy for in vivo stability, because degradation can differ across serum, plasma and whole blood^50^.

Overall, this study identifies position 5 as a useful but context-dependent optimisation site within two C5a-derived peptide scaffolds. The transferable feature was not a uniform improvement in every measured property, but the sensitivity of both scaffolds to hydrophobic modification at this position. The resulting analogues represent starting points for different optimisation objectives: BM1 P5M provided the most balanced pharmacological profile within the BM1 series, whereas BM221 A5Nle Abu6Ala showed improved serum stability without a corresponding improvement in C5aR1:C5aR2 functional discrimination. Future C5aR1 agonist design should balance productive target activation and pathway engagement against activity at C3aR and C5aR2, while separately considering proteolytic stability and other developability properties. More broadly, this study provides proof of principle that structure-guided computational workflows can focus peptide–GPCR optimisation on experimentally productive regions of sequence space, while demonstrating that multi-receptor and multi-pathway validation remains essential.

## Experimental Section

### Peptide Synthesis and Purification

Peptide synthesis was performed using Fmoc (9-fluorenylmethyloxycarbonyl)-based solid phase peptide synthesis (SPPS) on 2-chlorotrityl chloride resin on a 0.25 mmol scale. A ratio of 4 eq. of amino acid, 4 eq. of (2-(1*H*-benzotriazol-1-yl)-1,1,3,3-tetramethyluronium hexafluorophosphate (HBTU) and 8 eq. of *N,N*-diisopropylethylamine (DIPEA) was used for each coupling. N-terminal acetylation was performed on-resin using 2 × 5 min treatments of 5% acetic anhydride and 3% DIPEA in DMF at 25 °C. Following synthesis, the peptide was cleaved from the resin and sidechain protecting groups removed with a treatment of trifluoroacetic acid (TFA)/triisopropylsilane (TIPS)/water (95:2.5:2.5) for 2 hours. Following cleavage, purification was performed using reverse-phase high-performance liquid chromatography (RP-HPLC) using an increasing gradient of 1% buffer B (90% acetonitrile, 0.05% TFA) per minute in buffer A (0.05% TFA) over 80 minutes (Phenomenex Jupiter 300 Å, 10 µm, 250 × 21.2 mm).Analysis was performed using electrospray mass spectrometry (ESI-MS) (AB SCIEX API 2000) to identify fractions containing mass/charge ratios that matched the desired product. Purity was determined using analytical RP-HPLC (Agilent, 300 Å, 5 µm, 150 × 2.1 mm), with all peptides purified to >95% purity.

### Computational Modelling of C5aR1–Ligand Complexes

For each system, 100 models were generated, and those with the lowest discrete optimised protein energy (DOPE) scores were selected for further refinement. Selected models were energy minimized and subsequently subjected to molecular dynamics (MD) simulations in GROMACS 2018 using the AMBER99SB-ILDN force field. Energy minimisation was performed using 10,000 steps of the steepest descent algorithm, followed by an equilibration protocol in which positional restraints were gradually released. Restraints on the peptide were released first over 6 ns, followed by release of restraints on the receptor over 2.4 ns. Unrestrained production simulations were then performed for 100 ns. Long-range electrostatic interactions were treated using the particle mesh Ewald (PME) method with standard parameters. All bonds involving hydrogen atoms were constrained using the LINCS algorithm, allowing a 2 fs time step. Temperature was maintained at 27 °C using the velocity-rescale (V-rescale) thermostat, and pressure was maintained at 1 atm using isotropic Berendsen coupling. Short-range electrostatic and van der Waals interactions were truncated at 1.0 nm. Mutant peptide models were generated by substituting residue side chains using MODELLER v9.18, after which the same minimisation and MD simulation protocol was applied.

Unfolding free energies of wild-type and mutant peptide–receptor complexes were estimated using FoldX, which uses an empirically parameterised energy function derived from experimental data. Differences in unfolding free energy (ΔΔG) between mutant and wild-type complexes were used to infer changes in binding affinity. Values were classified as decreased affinity (ΔΔG ≥ 1 kcal/mol), similar affinity (−1 < ΔΔG < 1 kcal/mol), or increased affinity (ΔΔG ≤ −1 kcal/mol). Peptide analogues were designed based on predicted binding free energy changes.

### Cell Culture and In Vitro Signalling Assays

Human Embryonic Kidney-293 WT (HEK-293 WT) cells were maintained in DMEM supplemented with 10% fetal calf serum, 100 U/mL penicillin and 100 µg/mL streptomycin. Chinese hamster ovary cells stably expressing either C3aR (CHO-C3aR) or C5aR1 (CHO-C5aR1) were maintained in Ham’s F12 media supplemented with 10% FCS, 100 U/mL penicillin, 100 µg/mL streptomycin and 400 µg/mL G418. All cell lines were maintained in T175 flasks (37 °C, 5% CO2) and subcultured at 90% confluency using TrypLE Express. Recombinant expression levels for CHO-C3aR and CHO-C5aR1 are 27 ± 7.6 pmol/mg and 7.0 pmol/mg, respectively, based on the manufacturer’s certificate.

### BRET β-Arrestin Recruitment Assay

Ligand-induced β-arrestin recruitment was measured in HEK293 cells transiently transfected with C5aR1-Rluc8 and β-arrestin 2-Venus or C5aR2-Venus and β-arrestin 2-Rluc8 constructs using Lipofectamine 2000. 24 hours post-transfection, cells were dissociated using TrypLE Express and seeded onto white 96-well tissue culture plates (100,000 cells/well) in phenol red-free DMEM supplemented with 5% FBS. On the following day, cells were incubated with Nano-Glo Endurazine Live Cell Substrate for 2 hours (37 °C, 5% CO_2_). Emissions at 460–485 nm (Rluc8) and 520–545 nm (Venus) were measured on a Tecan Spark 20 M microplate reader for 5 reads, with an additional 15 reads following agonist addition. Ligand-induced BRET responses were calculated by dividing Venus emission by Rluc8 emission for each well. Responses were then normalised to the vehicle control and C5a control, which were defined as 0% and 100% response, respectively.

### pERK1/2 Signalling Assay

Ligand-induced phospho-ERK1/2 signalling was assessed using the AlphaLISA SureFire Ultra pERK1/2 (Thr202/Tyr204) assay kit according to the manufacturer’s instructions. CHO-C3aR or CHO-C5aR1 cells were seeded (50,000 cells/well) into 96-well tissue culture-treated plates, incubated for 24 hours and subsequently serum-starved overnight. Ligand dilutions were prepared in serum-free Ham’s F12 medium. Cells were stimulated with respective ligands for 10 minutes and then immediately lysed using AlphaLISA lysis buffer. Cell lysates (5 µL/well) were added to a 384-well ProxiPlate, followed by the donor and acceptor reaction mixes (2.5 µL/well each). Following a 2-hour incubation in the dark, the plate was read on a CLARIOstar Plus microplate reader following standard AlphaLISA settings.

### Serum Stability

Pooled human A/B serum was thawed on ice and clarified via centrifugation at 10,000 × g for 10 min, after which the supernatant was aliquoted into 1.5 mL microcentrifuge tubes and pre-equilibrated at 37 °C for 20 min alongside a water-only control. BM221, BM221 A^5^Nle, and BM221 A^5^Nle Abu^6^Ala were added to each tube to give a final concentration of 40 µM. At each time point, 40 μL of the reaction mixture was removed from each tube and combined with 40 μL of 6 M urea to denature serum proteins and halt proteolysis. Samples were incubated on ice for 10 min, after which 40 µL of 20% (w/v) trichloroacetic acid (TCA) was added, followed by a further 10-min incubation on ice to precipitate proteins. Quenched samples were centrifuged at 14,000 × g for 10 min at 4 °C, with 100 μL supernatant transferred to HPLC vials for further analysis. Intact peptide was quantified by RP-HPLC using the integrated peak area corresponding to the parent peptide. Serum stability was expressed as % peptide remaining, calculated by normalising the peak area at each time point to the corresponding t = 0 peak area for that peptide. Data represent mean ± SEM of n = 3 independent experiments.

### Data Collection and Analysis

Experiments were conducted in triplicate and conducted on at least three different days. Data were analysed using GraphPad Prism 10.1 and expressed as mean ± standard error of mean (SEM). For each repeat, data was normalised before being combined. Logarithmic concentration-response curves were plotted using combined data and analysed to calculate the potencies of each peptide. Receptor selectivity was quantified as the difference in potency between receptors, calculated as ΔpEC50 values for each peptide ligand.

#### Abbreviations

BRET: bioluminescence resonance energy transfer
C3aR: complement component 3a receptor
C5aR1: complement component 5a receptor 1
C5aR2: complement component 5a receptor 2
DOPE: discrete optimised protein energy
GPCR: G protein-coupled receptor
HPLC: high-performance liquid chromatography
MD: molecular dynamics
pERK: phosphorylated extracellular signal-regulated kinase
RP-HPLC: reversed-phase high-performance liquid chromatography
SAR: structure–activity relationship
SPPS: solid-phase peptide synthesis
TCA: trichloroacetic acid.

