## Supplemental Figures for "Structure-Guided Design of C5aR1-Selective Peptide Agonists"

### Suppporting Information

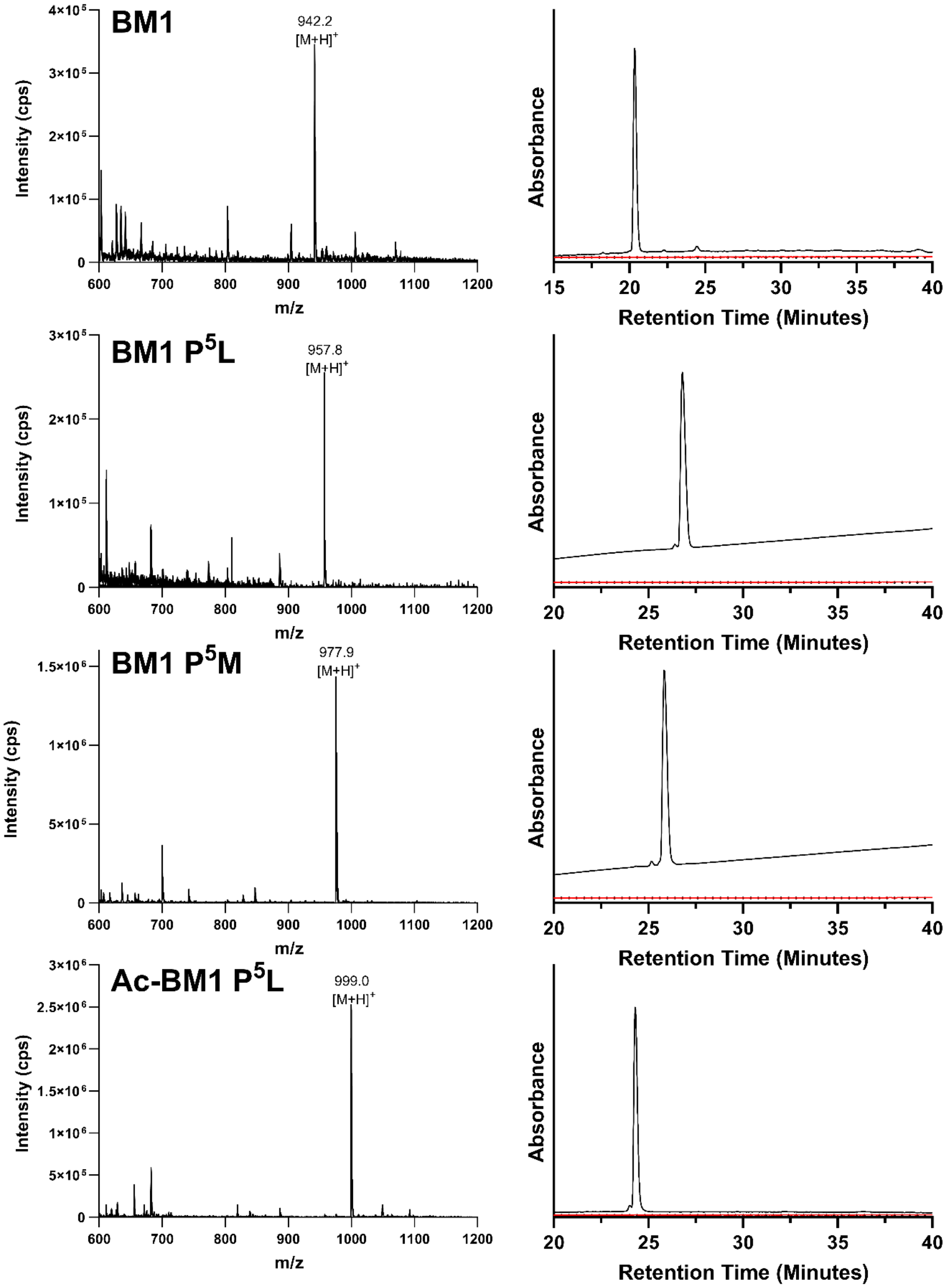


**Appendix 1:** Mass spectra and analytical HPLC traces of purified BM1 analogues. Analytical traces were run at an increasing gradient of 1% buffer B (90% ACN, 0.045% TFA) in buffer A (0.05% TFA) per minute using an Agilent, 300Å, 5 µm, 150 x 2.1 mm C18 column. Analysis was performed using electrospray mass spectrometry (ESI-MS) (AB SCIEX API 2000) to identify fractions containing mass-to-charge ratios that matched the desired product.

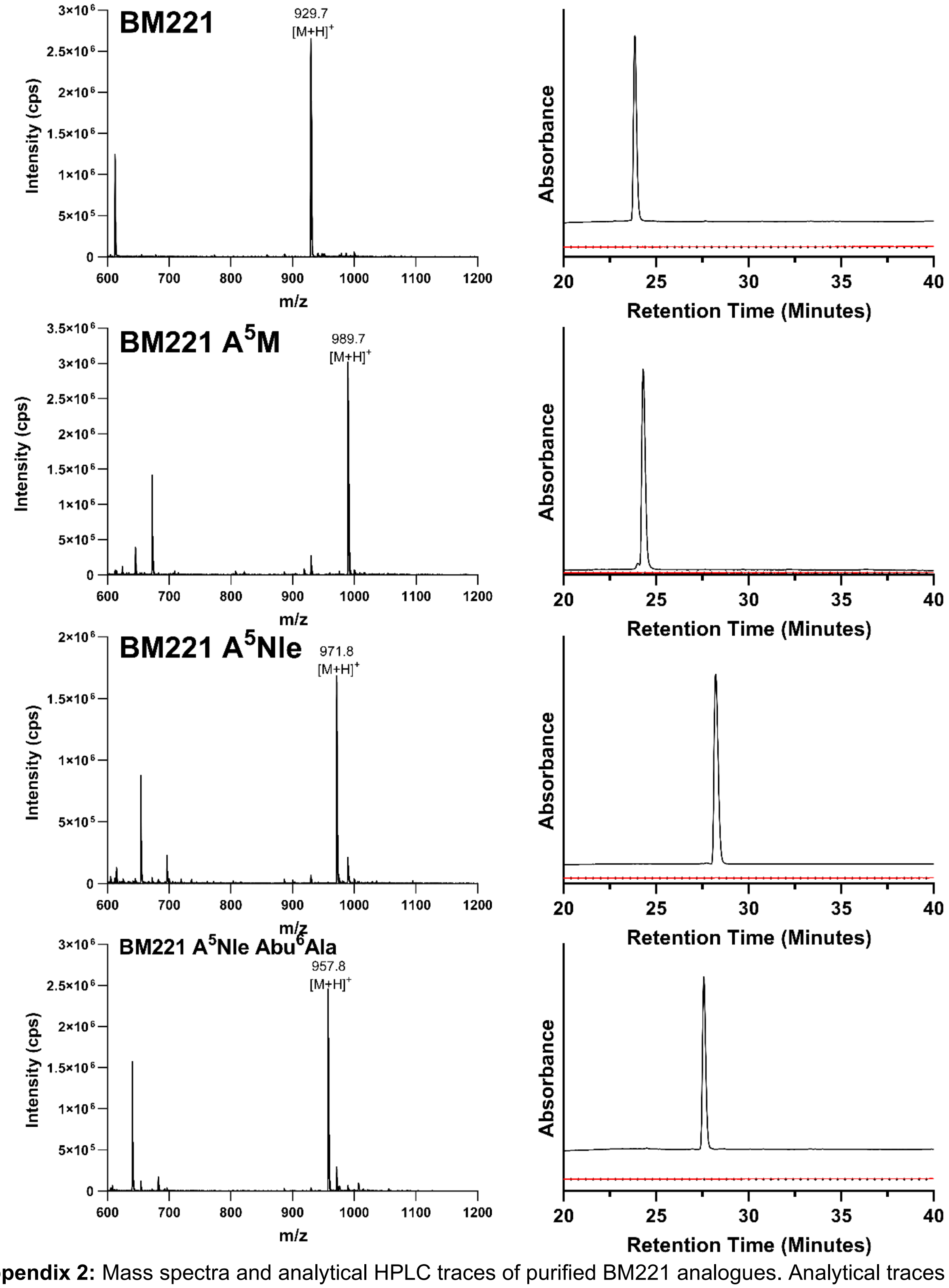


**Appendix 2:** Mass spectra and analytical HPLC traces of purified BM221 analogues. Analytical traces were run at an increasing gradient of 1% buffer B (90% ACN, 0.045% TFA) in buffer A (0.05% TFA) per minute using an Agilent, 300Å, 5 µm, 150 x 2.1 mm C18 column. Analysis was performed using electrospray mass spectrometry (ESI-MS) (AB SCIEX API 2000) to identify fractions containing mass-to-charge ratios that matched the desired product.
